# Linear alkylbenzenes (LABs) as sewage pollution indicators in the aquatic ecosystem of Peninsular Malaysia

**DOI:** 10.1101/863647

**Authors:** Sadeq. A. A. Alkhadher, Aeslina Abdul Kadir, Mohamad Pauzi Zakaria

## Abstract

Linear alkylbenzenes (LABs) as sewerage indicators were investigated in the sediments of the West and South Peninsular Malaysia. Surface sediment samples were analyzed using gas chromatography–mass spectrometry. Results showed that LAB concentration in the samples of Port Dickson and the Kim Kim River varied from 111.6 to 255.8 and from 88.2 to 119.0 ng·g^−1^ dry weight, respectively. The ratios of internal isomer, in which the benzene ring was close to the center of the linear alkyl chain, to external isomer, in which the benzene ring was near the end of the linear alkyl chain (I/E ratios), of LABs in Port Dickson coast sediments ranged from 2.6 to 4.1. By contrast, the ratios for sediment from the Kim Kim River varied from 1.7 to 1.9. I/E, long-chained LABs over short-chained LABs (L/S), and C_13_/C_12_ ratios indicated that the aquatic environment received primary and secondary sewage effluents. These findings emphasized the necessity of continued water treatment system development in Malaysia.

## Introduction

A significant habitat for organisms is the coastal environment. However, this sensitive environment has been severely impacted by anthropogenic activities and the rapid urbanization of the 20th century (Wei et al. 2014). The importance of coastal waters for recreation and aquaculture has been established (Law and Othman 1990); however, coastal areas are still commonly used as final disposal sites of human waste (Craig 2012).

Serious environmental pollution can be caused by industrial facilities and metropolitan centers along rivers that lead to the discharge of industrial and domestic wastewater, especially given inadequate regulation (Zhang et al. 2012). Surface water bodies have become polluted due to the uncontrolled release of sewage to the aquatic environment with partial or no wastewater treatment. Municipal wastewater to the aquatic environment has become a significant environmental issue in the last decades (Alkhadher et al. 2015). These effluents can result in the contamination of water, sediments, and aquatic species in addition to being detrimental to human health.

Linear alkylbenzenes (LABs) as organic pollutants are introduced to coastal and riverine environments accompanied by partially or nontreated municipal sewage and industrial effluent (Islam and Tanaka 2004; Oller et al. 2011). LABs with C_10_–C_14_ alkyl chains are primary materials for producing LAB sulfonate (LAS). LASs are manufactured via the sulfonation of LABs with H_2_SO_4_ or SO_3_ for use in synthetic detergents (Ricking et al. 2003). As a result of this incomplete sulfonation, LABs along with LAS detergents are released to the environment. These constituents are ubiquitous in many aquatic environments considering the widespread discharge of partially and nontreated municipal sewage to riverine and coastal sediments (Takada & Ishiwatari 1987; Wei et al. 2014; Dauner et al. 2015).

Sewage pollution can be assessed by chemical and microbiological markers (Vivian 1986; Takada and Eganhouse 1998). LABs are organic markers that are used for an anthropogenic indicator on the aquatic ecosystem given their exclusive association with human activities (Eganhouse 1997). Therefore, LABs are reported as chemical markers because of their widespread occurrence in aquatic environments known to be impacted by wastewater (Ishiwatari et al. 1983; Takada and Ishiwatari 1987; Hartmann et al. 2000).

Branched alkylbenzenes have completely changed to LABs for surfactant manufacture since the 1960s due to the superior rates of biodegradability and cost-effectiveness. Improved biodegradability of LABs relates to the position of phenyl-substitution groups on alkyl chains. For example, external isomers biodegrade faster than internal isomers. The isomeric structure and LAB concentration and their isomeric structure are due to the degree and type of wastewater released to the environment (e.g., raw sewage versus secondary sewage effluent), thereby making LABs ideal indicators of wastewater sources (Tsutsumi et al. 2002).

Peninsular Malaysia, especially along the west and south coasts, has undergone rapid industrialization and population over the last years. These developments have increased domestic and industrial wastewater contamination. Given frequent heavy precipitation in Malaysia, suspended particulate matter is commonly transported via surface runoff and flow into rivers; then, it undergoes sedimentation in estuaries (Shahbazi et al. 2010).

Contamination sources in the West and South Peninsular Malaysia must be monitored for health and environment because aquatic environments are used for recreation and fishing.

In Malaysia, the highest level of sewage contamination is shown in coastal areas where most of the country’s population live. A significant relationship occurs between sewage pollution and waterborne diseases in Asian countries (Isobe et al. 2002). Therefore, aquatic environment sediments and their potential sources must be investigated for possible sewage pollution to enhance water quality and decrease the danger of infectious illnesses (Wang et al. 2010).

The majority of the contaminants that enter the aquatic environment from anthropogenic sources eventually settle out of the water column via sedimentation (Abdullah et al. 1999). Therefore, sediments must be analyzed to evaluate the nature and extent of sewage pollution in the aquatic environment. The present study aims to evaluate sewage pollution at select coastal and riverine ecosystems along the west and south coasts of Peninsular Malaysia using LABs as molecular markers. This study also aims to investigate the LAB distribution and concentration in sediments of the selected areas in addition to their possible use as indicators of the extent of pollutant degradation in the coastal and riverine areas. Moreover, this study aims to assess the current efficiency of existing sewage treatment plants based on LAB isomer distributions in the selected areas.

## Methodology

### Study areas

The sediment samples were obtained from the Port Dickson Coast and the Kim Kim River of West and South Peninsular Malaysia, respectively (Figure 1). The sampling sites and the selected areas of Peninsular Malaysia are listed in Table 1.

**Figure 1.**
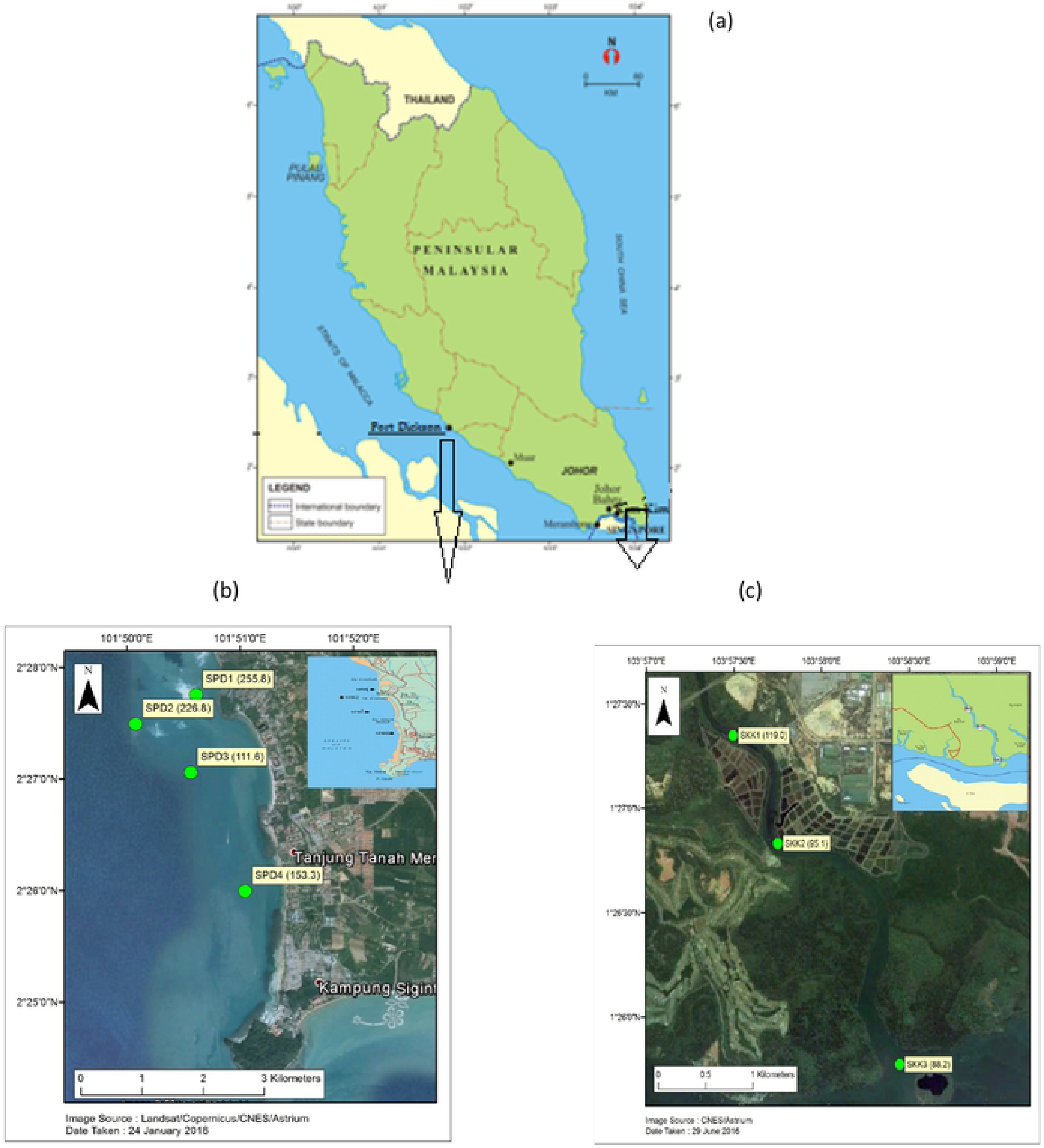
General location of study area, showing (a) a map of the Peninsular Malaysia, (b) Port Dickson Coast and (c) Kim Kim River.

### Sampling Process

The top four centimeters of sediment at select sample locations were sampled using an Ekman dredge. Surface sediments were moved to pre-cleaned aluminum foil and Ziploc bags and placed in a dried ice cooler box (Alkhadher et al. 2016). These sediments were then moved to a laboratory and kept out at −20 °C pending analysis.

### Analysis Process

The analysis process has been described by many researchers (Zakaria et al. 2002; Alkhadher et al. 2015). It includes homogenization, extraction, purification, and gas chromatography–mass spectrometry (GC-MS). Approximately 10 g of sample was weighed and then extracted in a Soxhlet glass extraction. A sample was spiked with 50 μL of 1-C_n_LAB mixture as the surrogate internal standard (SIS). Afterward, the sample was extracted with 300 mL dichloromethane (DCM) for 8–10 h. The elemental sulfur was removed by adding copper to the sample. Finally, the sample was reduced using a rotary evaporator.

The reduced sample was placed on top of the first-step silica gel column, which was packed with 5% H_2_O-deactivated silica gel (60–200 mesh size, Sigma Chemical Company, USA) to eliminate unwanted compounds. Hydrocarbons and nonpolar materials were eluted with 20 mL of hexane/DCM (3:1 v/v). The extract was then fractioned via second-step column chromatography with activated silica gel column.

The second hexane (4 mL) fraction was treated to obtain LAB fraction. The LAB fraction was reduced to nearly 1 mL and kept in a 2 mL amber vial. Consequently, it was further concentrated using N_2_. Then, the sample re-dissolved in 200 μL of isooctane includes 10 μg/g of internal injection standard. Finally, LABs were analyzed via GC-MS.

### GC-MS analysis of LABs

A total of 26 LAB isomers were analyzed using GC-MS detector. Aliquots (1 μL) of the purified sample were then transferred to the GC-MS injector. Helium was used as carrier gas (99.999% purity), with a controlled constant flow rate of 1.2 mL/min. The injection was performed at 70 °C for 2 min, with a ramp up of 30 °C/min until 150 °C. Temperatures were increased to 310 °C at a rate of 4 °C/min for 15 min. The selected ion monitoring (SIM) mode with splitless injection was performed in this analysis. LAB congeners were detected in the SIM mode at mass/charge ratios (m/z) of 91, 92, and 105.

LAB mixture standards were used as analytical standards. External calibration curves were illustrated with standard LAB mixtures for quantifying target LAB species. Concentrations of LAB species were determined by comparing retention times with the LAB mixture standards. Nondetectable species (<0.2 ng·g^−1^) were given zero concentration values in measuring the total LAB concentration.

### Quality control and assurance

This method was conducted via the sample analysis. Quality control analysis was performed by observing the SIS recovery before the extraction. The surrogate recovery was >83% of the samples. The sample with a recovery that is lower than the recognized value was reanalyzed. The LAB concentration was recovery-corrected against the surrogate standard (Cortazar et al. 2008). The LAB standards were prepared daily at the beginning of any analysis, and LAB blank samples were created in each batch. A blank process showed the free contamination of analytical system and glassware.

### Total organic carbon (TOC) analysis

Sediments were dried and homogenized using a pestle and a mortar. Then, the samples were acidified to remove the organic carbon content. The sample (2 g) was weighed, and two mL of 1 M HCl was added to the sample. The HCl was removed from the samples at 100 °C for 10 h. A LECO CR412 Carbon Analyzer (LECO Corporation, USA) at 1350 °C was used to determine TOC percentage (Nelson and Sommers 1996).

## Result and discussion

### Composition and Distribution

LABs varying in size from 5-phenyldecane to 2-phenyltetradecane and from C_10_ to C_14_ were investigated. A total of 26 LAB isomers were defined as n−C_m_ LAB, where *n* indicates the phenyl situation on the straight chain, and *m* denotes the alkyl carbon number. These isomers were expressed as a total sum (ΣLABs) and as a sum of LABs with C_10_–C_14_ (∑LABs C_10_–C_14_) in the collected sample (Table 2).

The composition of LAB isomers in the sediments of Port Dickson was measured (Figure 2). The C_13_ homolog species had the highest concentration, whereas the C_10_ concentration was lowest at locations SPD1, SPD2, SPD3, and SPD4.

**Figure 2.**
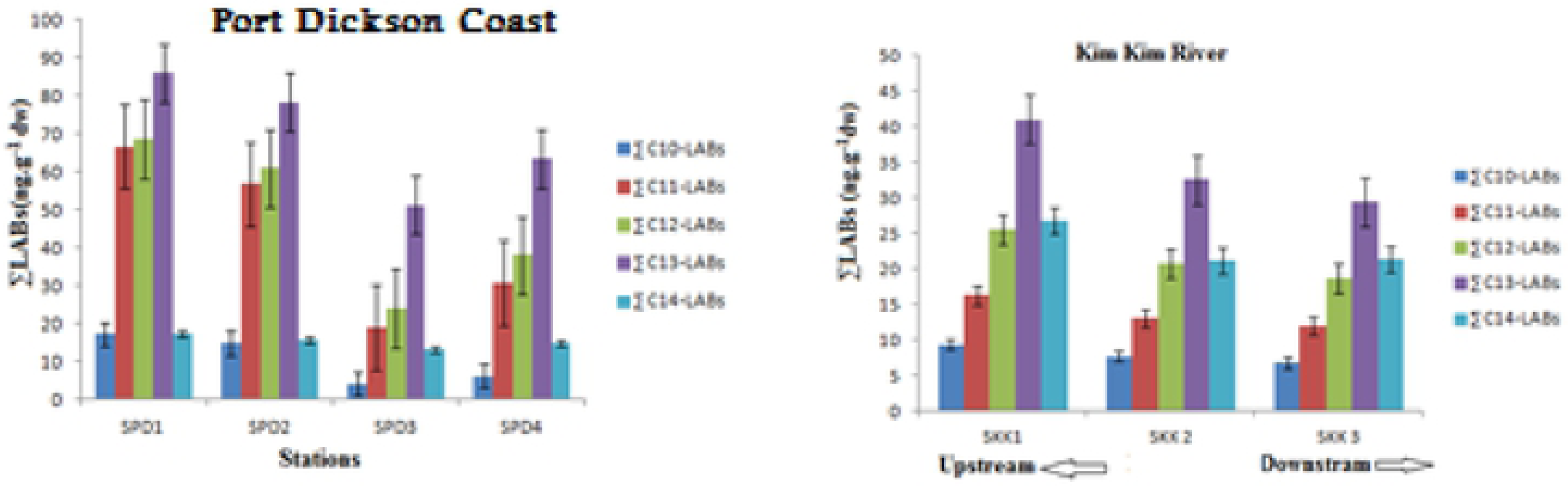
Compositional profiles of ∑LABs in Port Dickson Coast and Kim Kim River sediments

The compound analysis of ∑LABs in these surface sediments showed a high concentration of long-chain ∑LABs, such as tridecane (C_13_) and tetradecane (C_14_). Long-chain LABs (LC-LABs) in all Port Dickson samples showed higher concentrations than short-chain LAB compounds (SC-LABs). The SC-LABs, represented by the sum of decane (C_10_) and undecane (C_11_) homologs, showed the lowest concentration among the ∑LAB compounds in SPD1, SPD2, SPD3, and SPD4.

The isomeric compositions of ∑LABs are illustrated in Figure 2. The chain length of LAB distribution was dominant with C_13_ as the most abundant, followed by C_12_ and C_14_ homologs. C_10_ and C_11_ homologs were detected in low concentrations.

A high ∑LAB concentration with increasing chain length is expected given the high hydrophobicity of LABs (Sherblom et al. 1992). The chain length distributions of LABs in Port Dickson samples showed high concentrations of C_13_, followed by C_12_ and C_11_. C_14_ and C_10_ homologs were detected in low concentrations. Sedimentary distribution showed a decreased C_10_ homolog in comparison with those in sewage sludge and detergents (Luo et al. 2008).

The compositional profiles of isomeric and homologous LABs in sediments of the Kim Kim River were the highest for C_13_, followed by C_12_ and C_14_ homologs, as depicted in Figure 2.

Simultaneously, the lowest concentrations were observed for C_10_ and C_11_. These differences suggested the selective degradation of LABs with C_10_ and C_11_ homologs via the deposition of effluent particles. The isomeric composition in the sediments displayed the chain length distribution, with C_10_ and C_11_ homologs decreasing in comparison with those in sewage sludge and detergents (Luo *et* al. 2008).

The total ∑LAB concentration in this river was significantly controlled by LC-LABs. A compound-specific analysis of LABs showed that some compounds, such as 6-C_13_, 5-C_13_, and 6-C_12_ from LC-LABs, contributed to the high concentration in comparison with other isomers, thereby indicating the long-range transport of ∑LABs. The highest concentration of C_13_ was reported in sample SKK1. Elevated C_13_ concentrations indicated anaerobic degradation.

The surface sediments collected from this river have also indicated an increasing trend in ∑LAB concentrations during the last years (Isobe et al. 2004). In addition, LAB compound-specific analyses showed the elevated concentrations of some compounds, mostly LC-LABs, such as C_13_ (tridecane) and its alkyl substitute, in comparison with historical data (Isobe et al. 2004).

∑LAB concentrations in sediment samples collected from the study areas were lower than those measured in Malacca, Penang Estuary, Port Klang, Northern Tokyo Bay, Jakarta, and Kolkata (Takada et al. 1992; Isobe et al. 2004). In Table 4.2, ∑LABs were comparable with those in the Humber Estuary and Narragansett Bay (Raymundo and Preston 1992; Hartmann et al. 2000).

The ∑LAB concentrations determined by the present study have been presented to be higher than those in the South China Sea and Pearl River estuary (Luo et al. 2008). A ∑LABs comparison with those from Malaysia and the world emphasized that sewage pollution in the study areas is low to moderate.

∑LABs in the studied samples were distributed consistent with the following track: Port Dickson > Kim Kim River. On this basis, the geographical trend of sampling stations influenced ∑LAB distribution.

The findings of the current study showed that the Port Dickson sediment samples had a high level of ∑LABs. On this basis, high urbanization and industrialization in coastal areas may be the reason behind this spatial distribution (Shahbazi et al. 2010). Furthermore, the distributions of ∑LABs in the aquatic ecosystem are influenced by riverine runoff of sewage effluents and dilutions with natural organic matters (Zeng et al. 1997; Dauner et al. 2015).

### Evaluation of the TOC

LABs are known hydrophobic compounds with a strong tendency to attach to particulate organic matter once in the aquatic environment. As such, the TOC content in the sediments has been expected to be correlated with ∑LAB concentrations (Wang et al. 2001). TOC and water quality parameters were measured at each sampling location using proper equipment (Figure 4). TOC was measured in all sediments from the Port Dickson area and the relationship among ∑LABs, and TOC was assessed with linear regression and coefficient of determination (R^2^). The relationship was assessed using the R^2^ of 0.64. Therefore, the TOC may be a significant factor in spatial distributions of ∑LABs from the land to the studied stations at the Port Dickson coast. This result was consistent with the TOC results from the Dongjiang River, where ∑LAB concentrations were found to be linearly correlated with TOC concentrations (R^2^= 0.82), with wastewater discharge suggested as the main source for organic carbon in the sediments.

**Figure 3.**
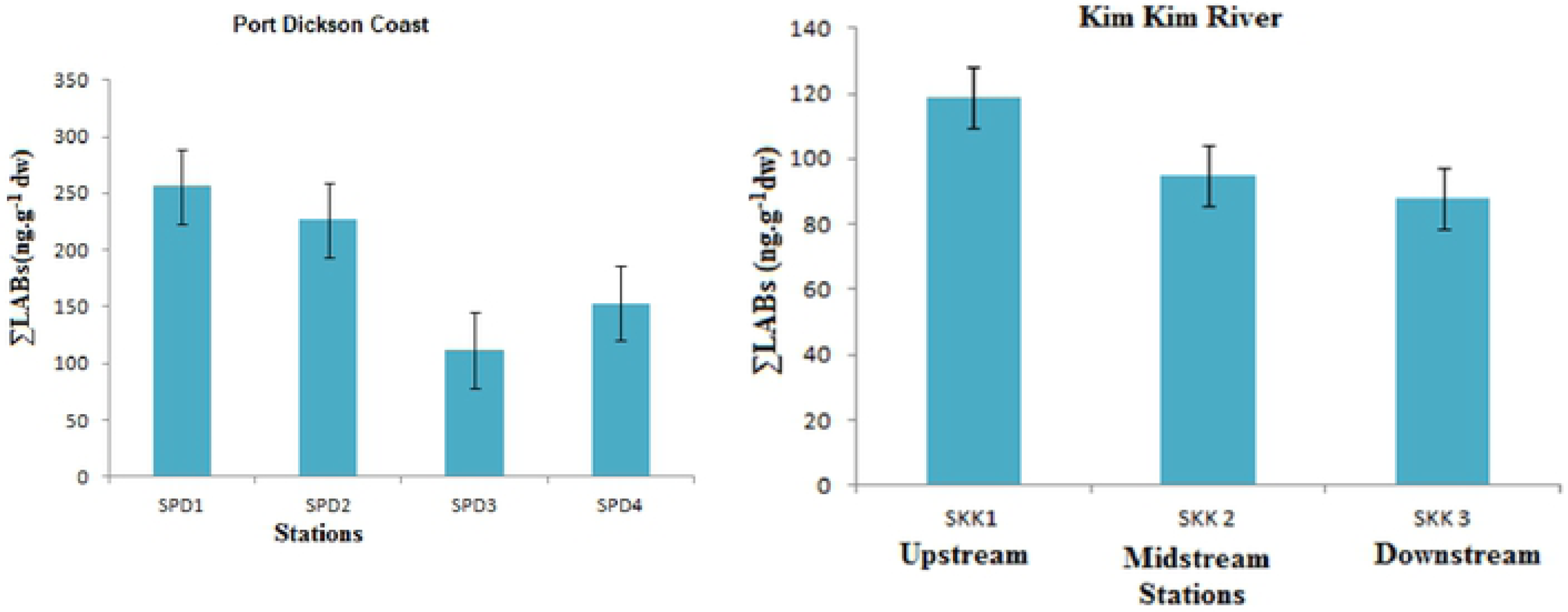
Concentration of ∑LABs in Port Dickson Samples and Kim Kim river, Standard error bars are shown

**Figure 4.**
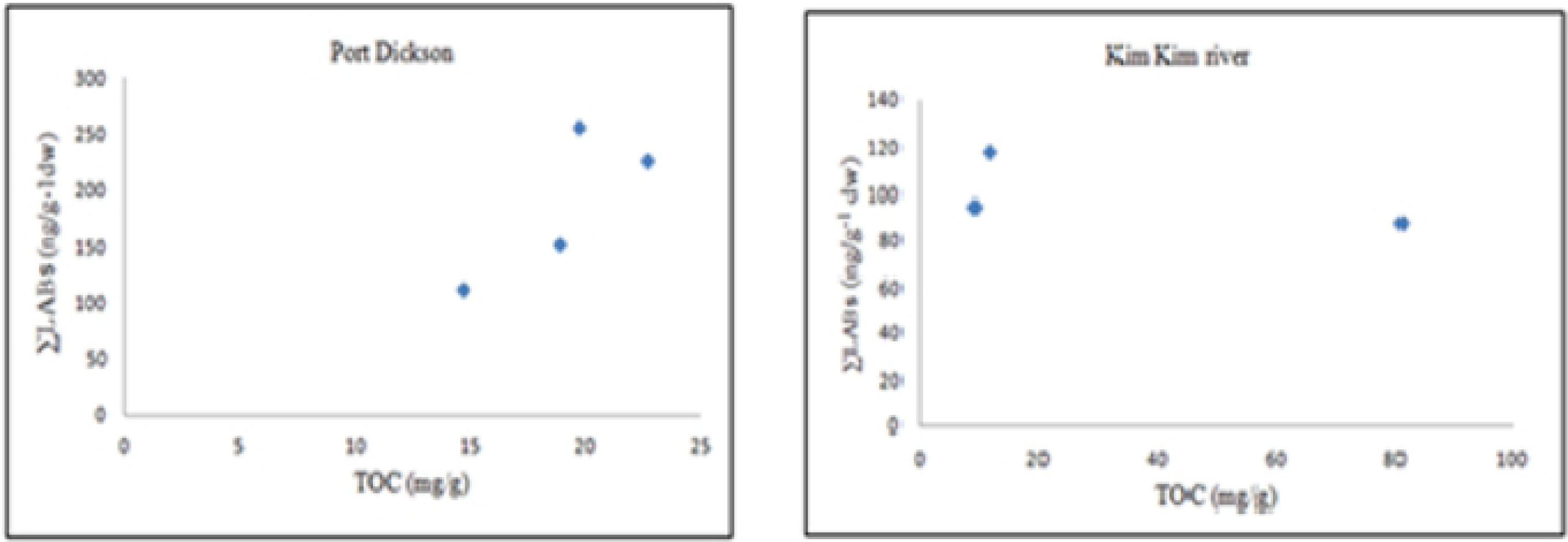
Scatter plots of ∑LABs and TOC in sediment samples of Port Dickson and Kim Kim river

The sediments of the Kim Kim River demonstrated a weak relationship between ∑LABs and TOC (R^2^=−0.42). Thus, TOC was not a determining factor in the distribution of ∑LABs in the sediments, and the intensity of a sewage input from different local sources was likely the key factor (Figure 4). Weak ∑LABs–TOC correlations determined for other areas of Peninsular Malaysia, such as the Selangor and Perak Rivers, were assessed using the R^2^ values of 0.008 and 0.17, correspondingly, with similar indications (Magam et al. 2015; Masood et al. 2015).

LAB degradation in surface sediments ranged from low in SKK3 from the Kim Kim River to high in SPD1 from the Port Dickson coast. The average ∑LAB biodegradation was 64% for the samples from the Port Dickson coast and 34% for the samples from the Kim Kim River. This result was attributed to greater selective biodegradation of ∑LAB species in the Port Dickson samples than in the Kim Kim River samples.

### Assessment of ∑LAB degradation and sewage treatment efficiency

The internal isomer to external isomer ratio (I/E ratio) results showed that most ∑LABs in Port Dickson come from secondary sewage effluents, whereas the dominant source of ∑LABs in the Kim Kim River was determined from primary sewage effluent. This result was attributed to the presence of more sewage treatment plants in Port Dickson than in the Kim Kim River (Table 2).

The I/E ratios of the Port Dickson station samples ranged from 2.0 to 4.1; on this basis, secondary sewage effluents were released (Figure 5). The I/E ratios are greater in the studied areas than in the Pearl River estuary (0.6–1.5) (Luo et al. 2008), thus indicating a large release of treated sewage effluents. The extent of ∑LAB degradation ranged from 40% to 64%, with a mean value of 53%, by assuming that biodegradation was under aerobic conditions (Takada and Ishiwatari 1990).

**Figure 5.**
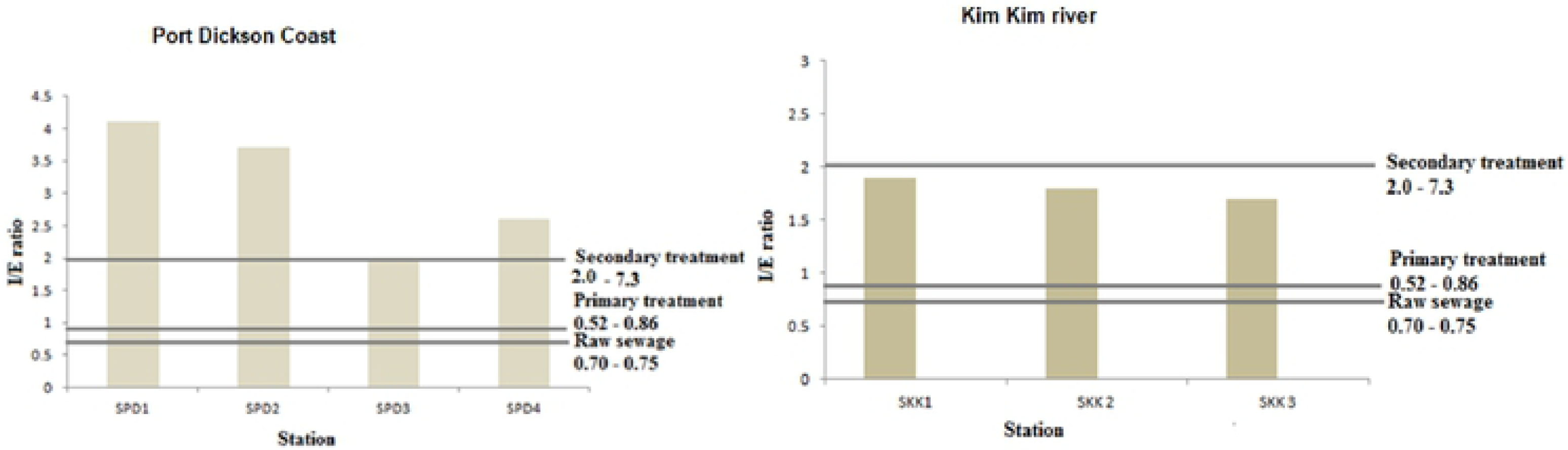
I/E ratios in sediments of Port Dickson Coast and KimKim River

LC-LAB/SC-LAB (L/S) ratios and C_13_/C_12_ were also calculated to confirm the extent of LAB degradation in the studied areas. As described previously, Port Dickson sediment is a receptor of well-traveled sewage discharge, and the ∑LAB signature partially supports this idea. A total of 82 sewage pipelines that release wastewater, including sewage from hotels and houses to the sea, lead to the deterioration of coastal water quality (Hamzah et al. 2011).

The I/E ratios of the Kim Kim River stations ranged between 1.7 at SKK 2 and 1.9 at SKK 1, with a mean value of 1.8; thus, primary treated sewage dominates inputs to the river water; this result is consistent with findings from Gustafsson et al. (2001).

The L/S ratios ranged from 2.4 to 2.6 with a mean value of 2.5, and they were higher than the 1.8 determined for detergents (Ni et al. 2008), thereby indicating the high biodegradation of ∑LABs in this area. By contrast, the ratio of C_13_/C_12_ ranged from 5.1 to 5.2, with a mean value of 5.14, which was higher than the 1.7 determined for coastal sediments (Liu et al. 2013).

The biodegradation values of ∑LABs were estimated, and they ranged from 34% to 38%, with an average of 36%. The direct release of waste from boats to the Kim Kim River in recent decades has been quite high because the river is used for recreational and fishing activities by locals (Abu Samah et al. 2011). Therefore, the direct release from riverboats has contributed to the occurrence of sedimentary ∑LABs, thereby accounting for the significant changes in molecular indices to some extent.

The ∑LAB inputs of developed countries were primarily from synthetic materials, whereas developing countries contributed less raw and primary sewage and more secondary inputs to the environment than the developed countries. Among the studies in surface sediment samples, the results revealed a primary effluent signature in the environment. ∑LABs that originated from different sources, such as municipal or industrial wastewater release, had a significant affinity for particulate matter, such as sewage particles and dissolved organic matter, thus possibly serving to enhance the transport of ∑LABs from their initial points of discharge via lateral movement (Harwood 2014; Zhang et al. 2012).

## Conclusions

The findings from the Port Dickson coast and the Kim Kim River sediment study showed that the LAB concentration ranged extensively. The results showed that Port Dickson coast is more contaminated by sewage effluents than the Kim Kim River. The LAB distribution in the study areas can be identified with sewage treatment degree in given output locations. The high level of LAB concentration observed in the study areas could be the result of inadequate sewage treatment plants that may not be afforded by the huge population in surrounding locations. Moreover, partially or nontreated sewage is being released to those areas given the overcapacity of treatment plants. Thus, the findings indicated that the sewage problem may continue as a result of increasing population. In the coming years, the magnitude of sewage released to coasts and rivers of Peninsular Malaysia in the future will be anticipated to increase broadly. The study findings also point the necessary actions that must be taken for wastewater treatment system enhancement. Thus, the incessant evaluation of current sewage pollution in coastal and riverine areas can be a means by which the improvement can be conducted by enhancing the current sewage system. Therefore, further investigation of sewage and other anthropogenic pollutants is required to decrease the health risk in coastal areas.

## Acknowledgments

The research was funded by the Inisiatif Putra Berkumpulan (IPB Grant no. 9412401). The authors are grateful to the chief editor and the reviewers of this article for their valuable contribution.

